# Proteomic profiling of Elp1-deficient trigeminal ganglia reveals disruption of neurotrophic and metabolic pathways in a familial dysautonomia mouse model

**DOI:** 10.64898/2025.12.05.692685

**Authors:** Carrie E. Leonard, Lauren Clarissa Tang, Beatrix Ueberheide, Lisa A. Taneyhill

## Abstract

**Background:** Elp1, a subunit of the Elongator complex, is essential for tRNA modification and neuronal development. Mutations in *ELP1* underlie familial dysautonomia (FD), a disorder marked by sensory and autonomic neuropathy. While loss of Elp1 disrupts trigeminal ganglion formation and survival, the downstream molecular consequences remain poorly defined.

**Results:** We performed quantitative proteomic profiling of trigeminal ganglia from Elp1 conditional knockout (CKO) and control embryos at E13.5. Across 5,650 detected proteins, 25 were significantly upregulated and 26 downregulated in Elp1 CKO embryos. EnrichR analysis revealed enrichment of upregulated proteins in amino acid transport and tRNA aminoacylation pathways, with links to neuromuscular and neuropathic diseases. Downregulated proteins were associated with RNA modification, cholesterol biosynthesis, and synaptic organization. Validation by immunohistochemistry confirmed decreased expression of the neurotrophic receptor Gfra3 and the neuropeptide Galanin, and increased levels of the chromatin regulator Chd1, in Elp1 CKO embryos relative to controls.

**Conclusions:** These findings demonstrate that Elp1 loss disrupts metabolic, RNA modification, and neurotrophic signaling pathways in the developing trigeminal ganglion. Proteomic analysis thus provides new insight into the molecular consequences of Elp1 deficiency and highlights candidate mechanisms contributing to sensory neuron vulnerability in FD.

## Introduction

The trigeminal ganglion, which mediates somatosensory input from the face to the brainstem, forms early in vertebrate development through the coordinated assembly of neural crest- and placode-derived precursors (D’Amico-Martel, 1982; D’Amico-Martel & Noden, 1983; Hamburger, 1961). While neural crest cells contribute both neurons and glia, placode cells produce only neurons in the trigeminal ganglion. Recent evidence suggests neural crest versus placodal progenitors preferentially give rise to distinct sensory neuron subtypes in the trigeminal ganglion, characterized by expression of tropomyosin receptor kinases (Trk), TrkA, TrkB, and TrkC (Leonard et al., 2022). Placode cells, which differentiate first, preferentially give rise to TrkB- and TrkC-expressing mechanoreceptors, while neural crest cells tend to become TrkA-expressing nociceptive neurons (Leonard et al., 2022). Together, these neuronal subpopulations work in concert to permit detection of sensations in the eye, forehead, nose, and upper and lower jaws.

Proper trigeminal ganglion development involves tightly regulated programs of neuronal specification, axon guidance, and target innervation, disruptions of which underlie a number of congenital sensory neuropathies (Moody & LaMantia, 2015; Park & Saint-Jeannet, 2010; Steventon et al., 2014; York et al., 2020). One such disorder is familial dysautonomia (FD), a rare autosomal recessive sensory and autonomic neuropathy caused by mutations in the *elongator acetyltransferase complex subunit 1 gene* (ELP1, also known as IKBKAP) (Anderson et al., 2001; Slaugenhaupt et al., 2001). FD patients have small trigeminal nerves and exhibit impaired facial pain and temperature perception, reflecting early defects in trigeminal sensory neuron development and survival (Geltzer et al., 1964; Gutiérrez et al., 2015; Mendoza-Santiesteban et al., 2017; Palma et al., 2018; Won et al., 2019). In a mouse model of FD where Elp1 is conditionally deleted in the Wnt1-Cre-expressing neural crest lineage (Elp1 CKO), trigeminal ganglion formation initiates normally but fails to progress due to defective axon pathfinding, target innervation, and subsequent loss of TrkA-expressing neurons (George et al., 2013; Leonard et al., 2022). While these findings underscore a requirement for Elp1 in the differentiation and maintenance of nociceptive neurons, other evidence suggests Elp1 is also required for the development of placode-derived neurons in the trigeminal ganglion and other cranial ganglia (Hines & Taneyhill, 2025; Tolman et al., 2022).

Elp1 encodes a scaffolding subunit of the six-member dimeric Elongator complex, initially characterized for its role in transcriptional elongation, but now known to be essential for the post-transcriptional modification of tRNAs and efficient translation of codon-biased transcripts (Esberg et al., 2006; Goffena et al., 2018; Huang et al., 2005, 2008; Karlsborn, Tükenmez, Mahmud, et al., 2014; Xu et al., 2015). In animal and *in vitro* models, Elp1 deficiency is also associated with impaired neurotrophic signaling, cytoskeletal abnormalities, mitochondrial dysfunction, and changes in gene expression (Cameron et al., 2021; Y.-T. Chen et al., 2009; George et al., 2013; Ghosh et al., 2021; Goffena et al., 2018; Jackson et al., 2014; Johansen et al., 2008; Ueki et al., 2018). Despite these central functions in protein homeostasis and cell signaling, the molecular mechanisms linking Elp1 loss to neurodevelopmental disruption are not well understood. Notably, most studies to date have focused on transcriptional consequences of Elp1 loss, yet proteomic changes may more directly reflect the cellular dysfunction that drives neuronal loss.

To address this gap, we performed a proteomics-based screen to identify differentially expressed proteins in E13.5 trigeminal ganglia from Elp1 CKO and littermate control embryos. This developmental stage coincides with the onset of TrkA neuron loss and disrupted innervation patterns observed in Elp1 CKO embryos, providing a critical window to capture proteomic alterations associated with early neurodevelopmental dysfunction. Our analysis reveals changes in proteins involved in cholesterol biosynthesis, growth factor signaling, and amino acid transport, offering new insight into the molecular underpinnings of sensory neuron vulnerability in FD.

## Results

To examine which proteins are differentially expressed in the Elp1 conditional knockout (CKO) compared to control samples, we dissected trigeminal ganglia from embryonic day (E)13.5 embryos and subjected them to mass spectrometry analysis (n=5 control, n=5 Elp1 CKO). Proteomic profiling of E13.5 trigeminal ganglia identified a total of 5,650 proteins across Elp1 CKO and control samples. Among these, 25 proteins were significantly upregulated and 26 were significantly downregulated in the Elp1 CKO ganglia compared to controls using a p-value <0.05 adjusted for multiple hypothesis testing (Benjamini & Hochberg, 1995). Of those, Elongator subunits Elp1, Elp2, and Elp3 were all significantly downregulated with Log2 fold changes of −1.706 (p_adj_=0.0005), −0.774 (p_adj_=0.0191), and −0.978 (p_adj_=0.0052), respectively (Supplemental Table 1).

EnrichR analysis (E. Y. Chen et al., 2013; Kuleshov et al., 2016; Xie et al., 2021) of the upregulated set of proteins highlighted potential transcriptional regulation by Myc and Runx1 (Table 1). Gene Ontology (GO) Biological Process categories were enriched for amino acid metabolism and transport, including “sulfur amino acid transport”, “tRNA aminoacylation for protein translation”, “amino acid import across the plasma membrane”, “L-amino acid transport”, and “neutral amino acid transport” (Table 1). Jensen Diseases terms associated with the upregulated proteins included several neuromuscular and neuropathic conditions, such as Charcot-Marie-Tooth disease, neuromuscular disease, trichothiodystrophy, spinal muscular atrophy, and mitochondrial myopathy (Table 1). Consistent with these findings, Reactome Pathways analysis identified significant enrichment in “cytosolic tRNA aminoacylation”, “tRNA aminoacylation”, “amino acid transport across the plasma membrane”, and broader categories of metabolism and metabolism of amino acids and derivatives (Table 1).

**Table 1.**
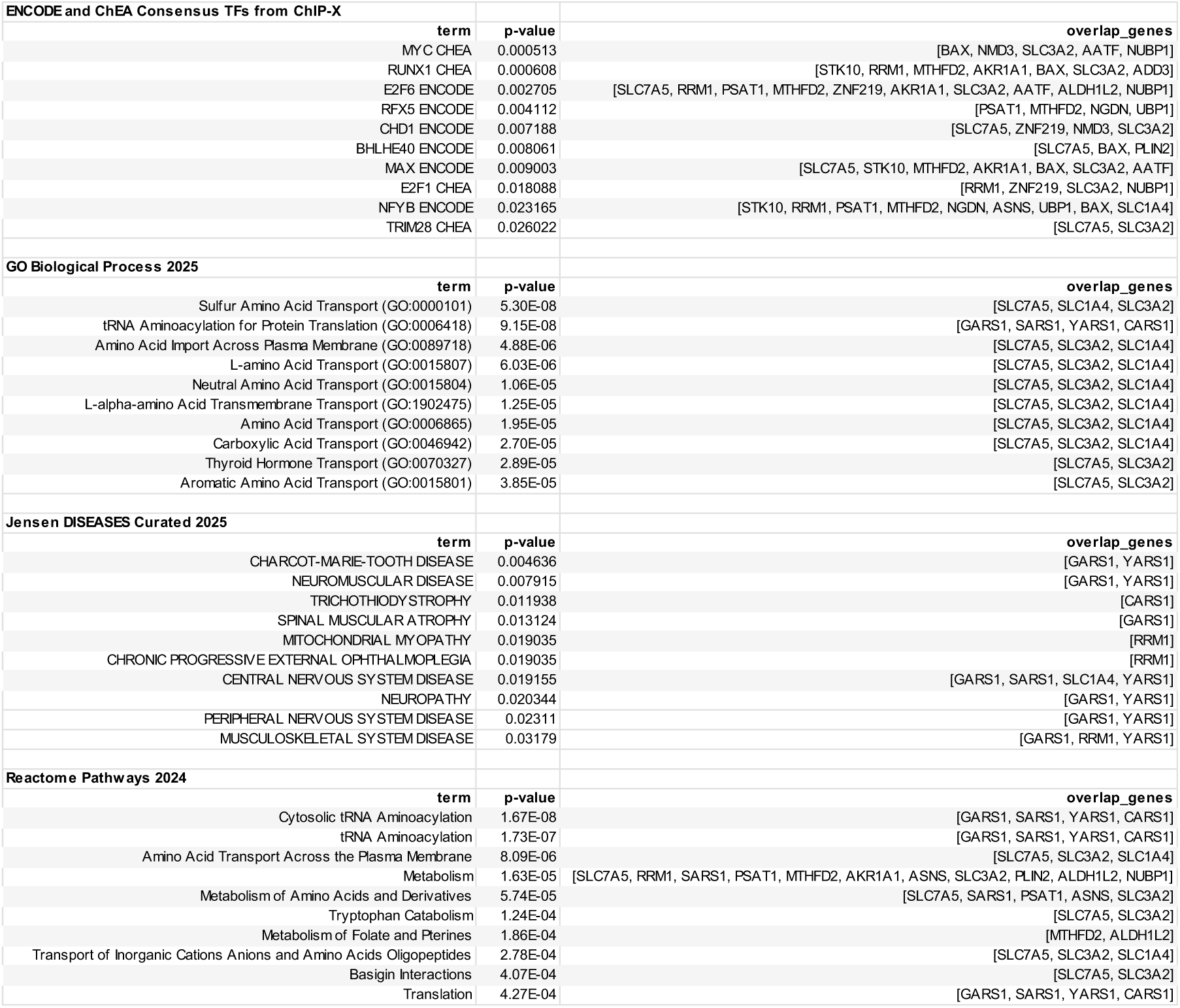
Top significant terms associated with proteins upregulated in E13.5 Elp1 CKO compared to control embryos, as identified by EnrichR analysis.

In contrast, the 26 downregulated proteins were associated with a distinct set of transcription factors, including KLF4, TCF3, CREB1, SP2, and SPI1 (Table 2). GO Biological Process terms highlighted processes related to RNA modification and synaptic regulation, such as “tRNA wobble base modification”, “tRNA wobble uridine modification”, “regulation of postsynapse organization”, “mRNA catabolic process”, and “regulation of translation” (Table 2). Jensen Diseases categories included hereditary sensory neuropathy, as well as opiate dependence and substance dependence (Table 2). Reactome Pathways analysis revealed significant enrichment in “cholesterol biosynthesis”, “activation of gene expression by SREBF”, “regulation of cholesterol biosynthesis by SREBF”, and “histone acetylation by HATs” (Table 2).

**Table 2.**
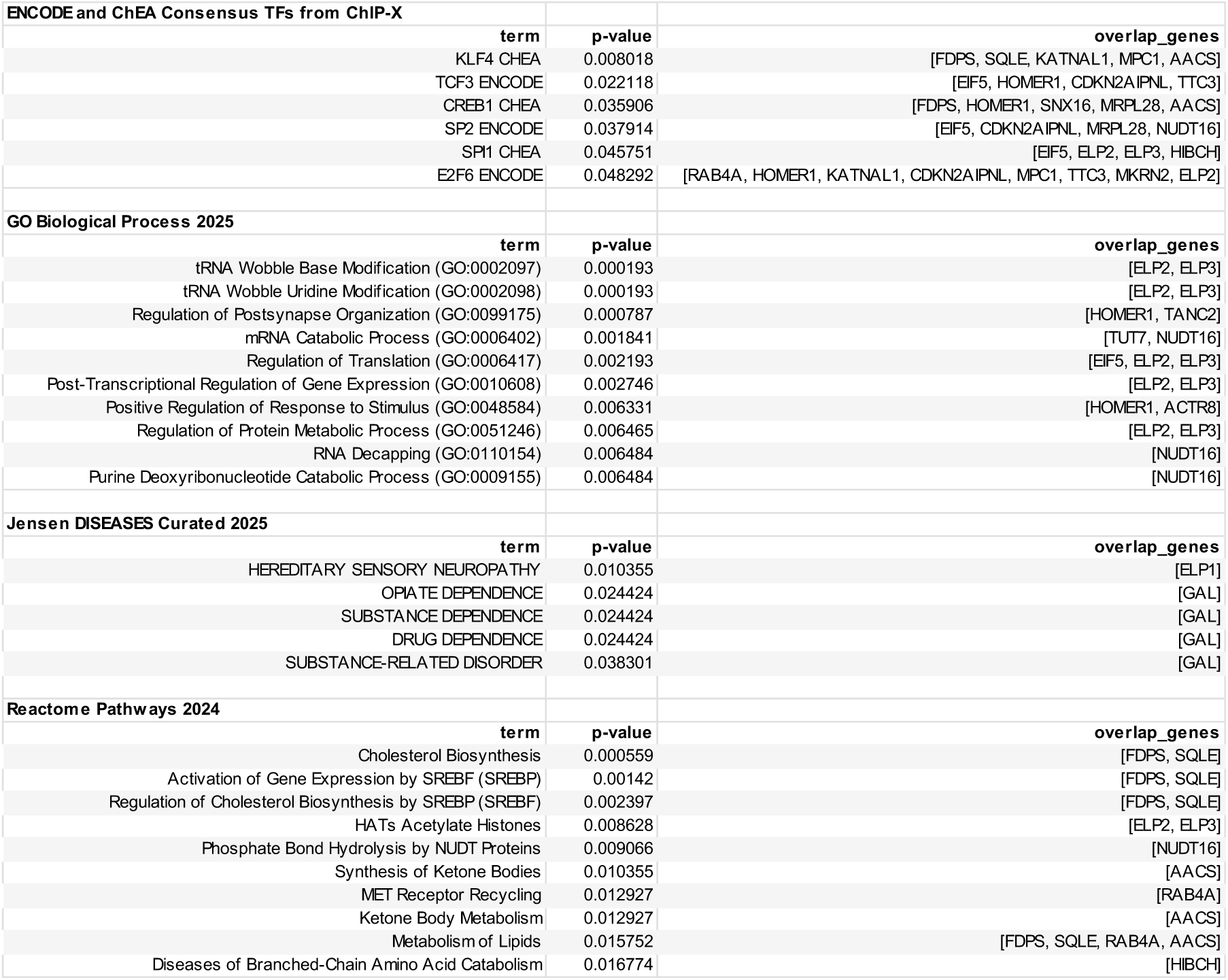
Top significant terms associated with proteins downregulated in E13.5 Elp1 CKO compared to control embryos, as identified by EnrichR analysis.

To validate the findings from mass spectrometry analysis, immunohistochemistry was performed to visualize select candidate proteins, including the GDNF family receptor alpha 3 (Gfra3), the neuropeptide Galanin (Gal), and Chromatin Helicase DNA-binding Protein 1 (Chd1). Data from the Allen Developing Brain Atlas reveals that Gfra3 is normally highly expressed in the trigeminal ganglia relative to surrounding tissues (Figure 1A,Allen Institute for Brain Science, 2024)). In the proteomics study, Gfra3 decreased by a Log2 fold change of −0.541 (p_adj_=0.0345) in the Elp1 CKO compared to controls (Supplemental Table 1). Gfra3 was confirmed via immunohistochemistry to be downregulated in the Elp1 CKO trigeminal ganglia compared to control trigeminal ganglia, with mean fluorescent intensity measuring 6477 relative fluorescence units (RFU) versus 8793 RFU, respectively (p=0.0006) (Figure 1B-D). Similar to Gfra3, Gal is normally observed in the trigeminal ganglia at higher levels than surrounding tissues (Figure 2A, (Allen Institute for Brain Science, 2024)). In the Elp1 CKO, Gal was decreased by a Log2 fold change of −1.285 compared to control (p_adj_=0.0021, Supplemental Table 1). Immunohistochemistry demonstrated that Gal was expressed at lower levels in the Elp1 CKO trigeminal ganglia, with a mean fluorescent intensity of 1501 RFU compared to 1784 RFU in control trigeminal ganglia (p=0.0038), further confirming data from the proteomic analysis (Figure 2B-D). Finally, Chd1 was upregulated with a Log2 fold change of 0.836 in Elp1 CKO compared to control samples (p_adj_=0.0379, Supplemental Table 1). Chd1 was also measured at higher levels via immunohistochemistry in the Elp1 CKO compared to controls, with mean intensity fluorescence of 14511 RFU versus 13131 RFU, respectively, in the trigeminal ganglia (p=0.0330, Figure 3). There was no data available from the Allen Developing Brain Atlas to demonstrate the normal expression of Chd1 in the embryonic trigeminal ganglion. Altogether, these findings confirm that candidate expression profiles via immunohistochemistry mirror results from the proteomic analysis, demonstrating that key proteins are significantly differentially expressed in the Elp1 CKO trigeminal ganglion compared to controls.

**Figure 1.**
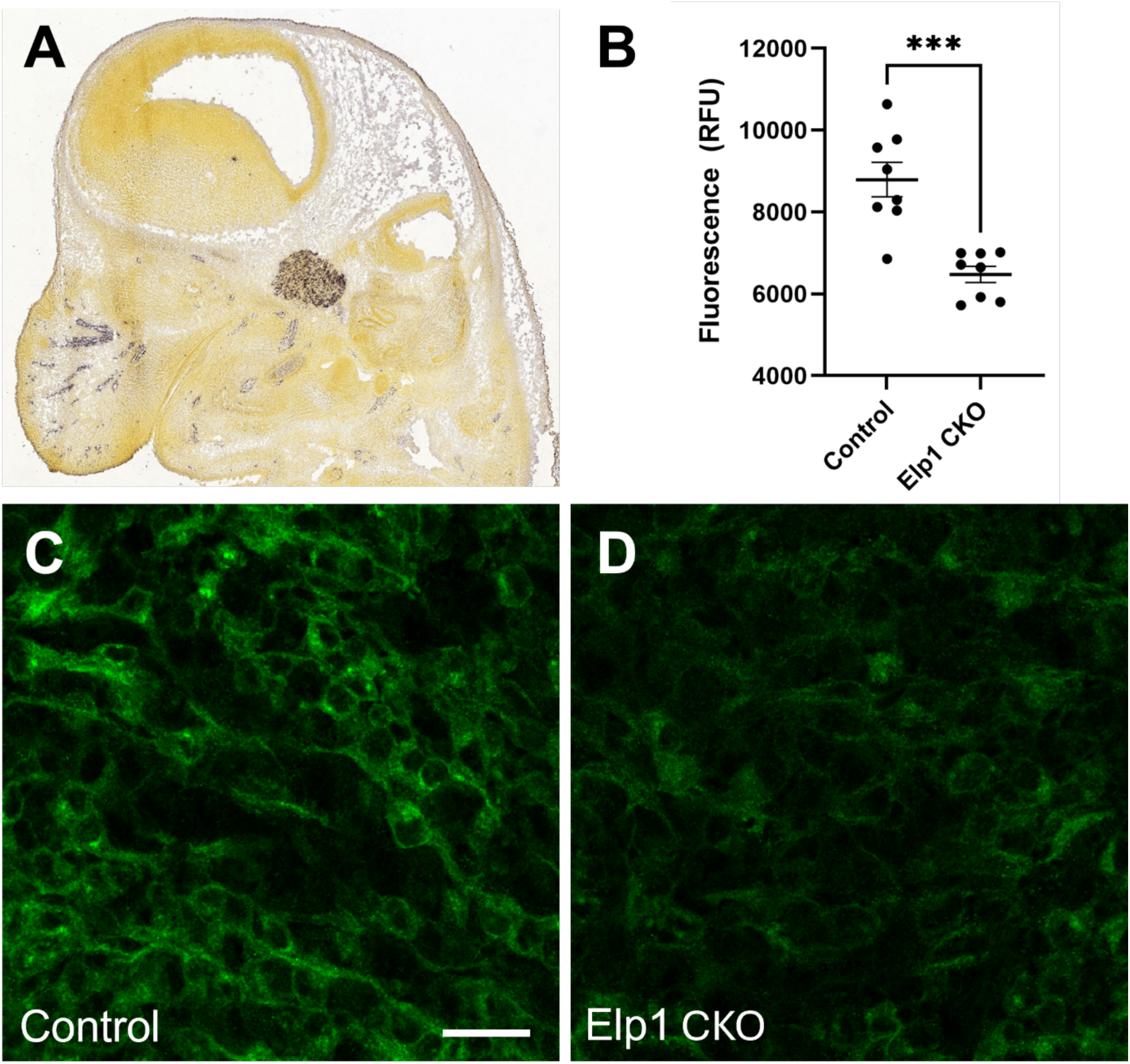
Gfra3 protein levels are decreased in E13.5 Elp1 CKO compared to controls. **A**, *In situ* hybridization data from the Allen Developing Brain Atlas showing expression of *Gfra3* mRNA in the E13.5 trigeminal ganglion (https://developingmouse.brain-map.org/experiment/show/100041840). **B**, Quantification of mean relative fluorescence units (RFU) of immunohistochemistry signal for Gfra3 in E13.5 control (**C**) or Elp1 CKO (**D**) trigeminal ganglia, ***p=0.0006, Welch’s t test. Scale bar in C equals 20µm and also applies to D.

**Figure 2.**
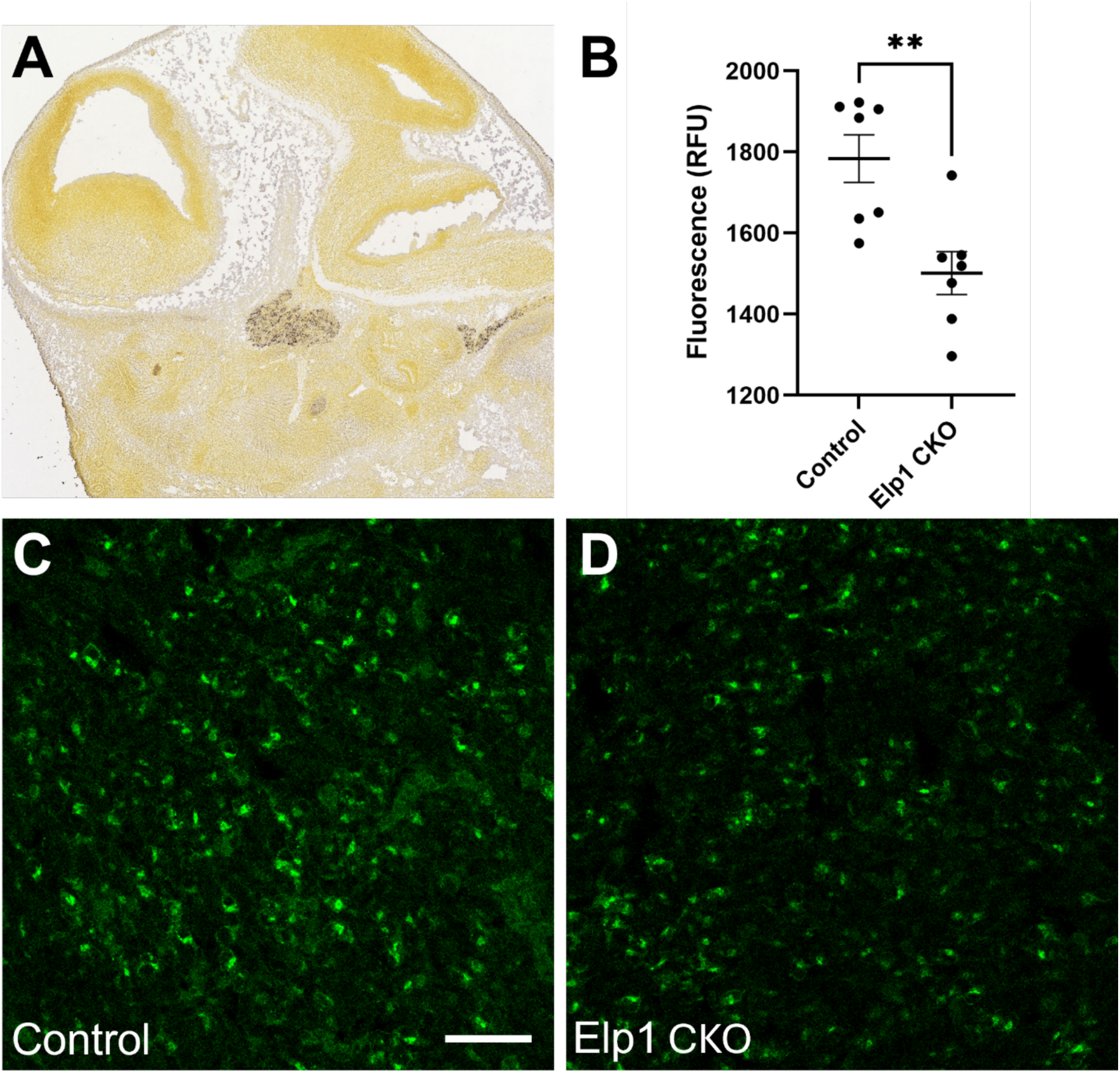
Gal protein levels are decreased in E13.5 Elp1 CKO compared to controls. **A**, *In situ hybridization* data from the Allen Developing Brain Atlas showing expression of *Gal* mRNA in the E13.5 trigeminal ganglion (https://developingmouse.brain-map.org/experiment/show/100046275). **B**, Quantification of mean relative fluorescence units (RFU) of immunohistochemistry signal for Gal in E13.5 control (**C**) or Elp1 CKO (**D**) trigeminal ganglia, **p=0.0038, Welch’s t test. Scale bar in C equals 50µm and also applies to D.

**Figure 3.**
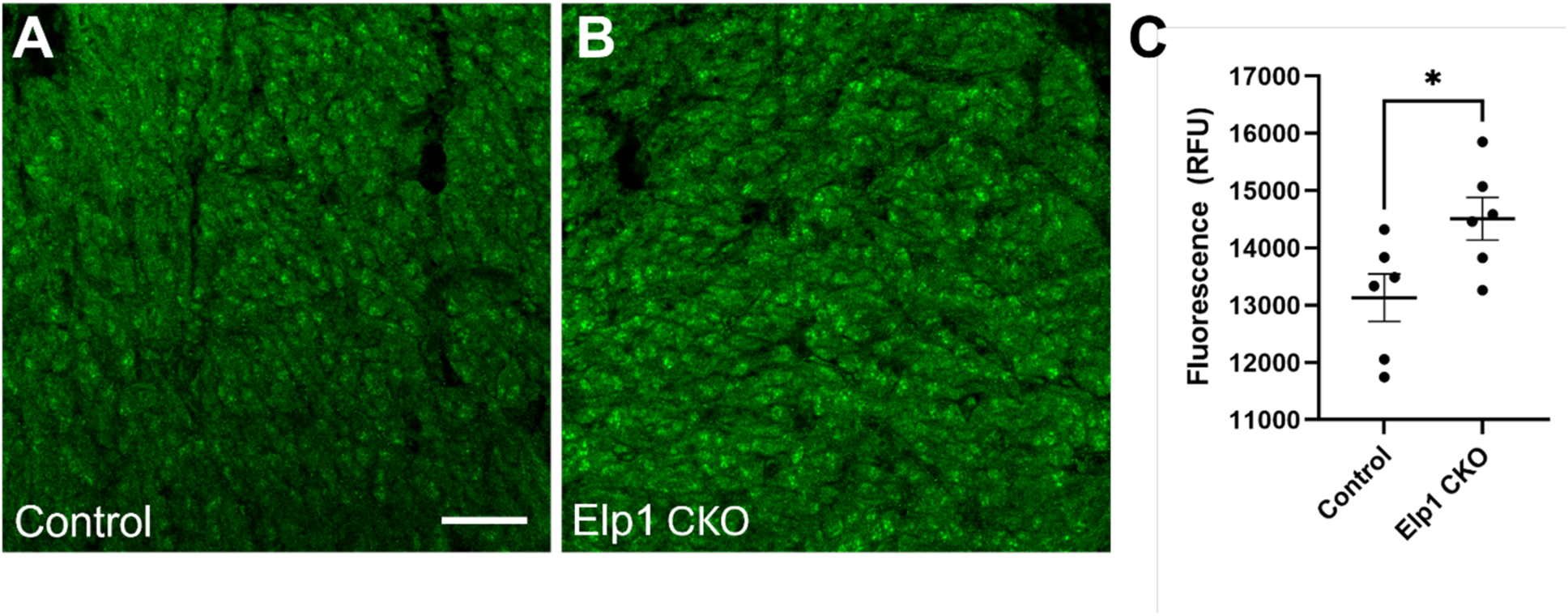
Chd1 protein levels are increased in E13.5 Elp1 CKO compared to controls. **A-B**, Immunohistochemistry showing expression of Chd1 protein in E13.5 control (A) or Elp1 CKO (**B**) trigeminal ganglion. **C**, Quantification of mean relative fluorescence units (RFU) of immunohistochemistry signal for Chd1 in E13.5 control or Elp1 CKO trigeminal ganglia, *p=0.0330, Welch’s t test. Scale bar in A equals 20µm and also applies to B.

## Discussion

Our proteomic analysis of E13.5 trigeminal ganglia from Elp1 CKO embryos provides new insight into the molecular consequences of Elp1 loss during cranial sensory neuron development. Although relatively few proteins reached statistical significance, both upregulated and downregulated sets were enriched for biological processes and disease terms directly relevant to neuronal metabolism, survival, and function. These results expand upon prior transcriptomic and histological findings (Goffena et al., 2018; Harripaul et al., 2024; Leonard et al., 2022), underscoring the essential role of Elp1 in maintaining the molecular integrity of developing trigeminal ganglia.

EnrichR analysis (E. Y. Chen et al., 2013; Kuleshov et al., 2016; Xie et al., 2021) pointed to possible involvement of transcriptional regulators such as Myc and Runx1 (associated with Elp1 CKO upregulated proteins, Table 1) and KLF4, TCF3, and CREB1 (associated with Elp1 CKO downregulated proteins, Table 2). Notably, these transcription factors were not themselves detected as differentially expressed in Elp1 CKO embryos (Supplemental Table 1), suggesting that altered activity or localization of the transcription factors may instead reflect changes in their downstream targets. Many of these regulators have well-established functions in cell growth, neuronal development, and plasticity (Jha et al., 2023; Kobayashi et al., 2012; Qin et al., 2011; Sakamoto et al., 2010), raising the possibility that the loss of Elp1 perturbs transcriptional networks required for balancing progenitor maintenance, growth, and division with neuronal differentiation. Such dysregulation could contribute to the cellular imbalances previously observed in Elp1 CKO trigeminal ganglia, including premature neuronal loss and disrupted axon outgrowth (Leonard et al., 2022).

The GO Biological Process categories identified among upregulated proteins highlight metabolic pathways, including amino acid transport and tRNA aminoacylation, processes essential for protein synthesis and neuronal survival. Alterations in amino acid metabolism have been linked to neuronal dysfunction in multiple neurodevelopmental and neurodegenerative contexts (Adla et al., 2024; He et al., 2025; Maszka et al., 2023), raising the possibility that metabolic stress contributes to the progressive loss of Elp1-deficient neurons. Conversely, downregulated proteins were enriched for pathways related to RNA modification, including tRNA wobble base modification. This result is consistent with the known function of Elp1, Elp2, and Elp3 as components of the Elongator complex, which mediates tRNA modifications necessary for accurate translation(C. Chen et al., 2009; Huang et al., 2005; Karlsborn, Tükenmez, Chen, et al., 2014; Karlsborn, Tükenmez, Mahmud, et al., 2014). The coordinated downregulation of Elongator subunits underscores the central role of this complex in the observed phenotype and provides a direct molecular connection to the pathogenesis of FD, where Elp1 mutations compromise neuronal function through impaired gene expression, in addition to other cellular processes. However, it is also possible that individual Elp1, Elp2, or Elp3 subunits independent of the Elongator complex may play unique roles in the dysregulation of molecular pathways leading to FD phenotypes.

Candidate protein validation further supports the link between Elp1 loss and sensory neuron dysfunction. Gfra3, which was significantly reduced in Elp1 CKO trigeminal ganglia, is predominantly expressed in nociceptive sensory neurons during development (Orozco et al., 2001) and mediates signaling by the neurotrophic factor Artemin (Baloh et al., 1998; Naveilhan et al., 1998). Reduced Gfra3 expression therefore suggests compromised trophic support for nociceptors, consistent with the depletion of TrkA associated with Elp1 loss in trigeminal neurons (Leonard et al., 2022). Similarly, Galanin, which was downregulated in the Elp1 CKO, has dual roles in supporting sensory neuron survival and modulating pain signaling in a context-dependent manner (Fonseca-Rodrigues et al., 2022; Holmes et al., 2000; Zhang et al., 1998). Its regulation by neurotrophic signaling, including NGF, places Galanin within a broader network of trophic factor–dependent mechanisms disrupted by Elp1 deficiency (Liu & Li, 2009; Shadiack et al., 1998). In contrast, Chd1 was upregulated in Elp1 CKO embryos. Notably, Chd1 was also indicated as a transcriptional regulator of upregulated proteins in the Elp1 CKO, according to EnrichR analysis (Table 1). Chd1 has established roles in positively regulating chromatin accessibility, stem cell pluripotency, and sensory perception (Bulut-Karslioglu et al., 2021; Gaspar-Maia et al., 2009; Schoberleitner, 2022), raising the possibility that its elevation could induce a more progenitor-like state in Elp1-deficient neurons. Collectively, these candidate validations highlight how altered expression of individual genes can provide mechanistic links between Elp1 loss, disrupted neurotrophic signaling, and sensory neuron vulnerability.

Several limitations of the present study should be acknowledged. First, the magnitude of protein expression changes was modest, even for Elp1 itself, likely reflecting the mosaic nature of the Elp1 CKO. Because Elp1 deletion was restricted to neural crest–derived lineages (George et al., 2013), placode-derived neurons within the trigeminal ganglion retain Elp1 expression, diluting the detectable effects at the whole-tissue level (Leonard et al., 2022). Furthermore, the trigeminal ganglion contains both neurons and glia, and our bulk proteomics approach does not allow cell type–specific resolution. While expression patterns of some proteins (e.g., Gfra3, Galanin) suggest neuronal origins based on their function and expression patterns, future studies employing cell type–specific isolation or single-cell proteomics will be required to discern neuronal versus glial contributions. Finally, our analysis was restricted to E13.5, a stage at which apoptosis and axonal abnormalities are already evident in Elp1 CKO ganglia(Leonard et al., 2022). Earlier developmental stages may reveal primary molecular disruptions preceding cell death, while later stages could clarify how persistent metabolic, transcriptional, and/or translational changes contribute to disease progression.

In conclusion, our proteomic study identifies discrete yet functionally significant changes in protein expression within the Elp1 CKO trigeminal ganglion. Altered pathways highlight disruptions in amino acid metabolism, RNA modification, and neurotrophic signaling, processes that are critical for neuronal survival and consistent with the loss of sensory function in FD. By validating key candidate proteins, this work establishes a framework for future studies aimed at dissecting cell type–specific mechanisms and exploring therapeutic interventions that restore metabolic and trophic support to vulnerable sensory neurons.

## Experimental Procedures

### Tissue Collection and Processing

Timed pregnant females were euthanized by CO₂ asphyxiation, followed by cervical dislocation to ensure death. Embryos were dissected and immediately placed into chilled 1× phosphate-buffered saline (PBS). For genotyping purposes, one hindlimb bud was harvested from each embryo. Whole embryos were fixed by immersion in 4% paraformaldehyde (PFA) in 1× PBS with gentle agitation at room temperature for two hours. Fixed specimens were rinsed three times in 1× PBS for 20 minutes per rinse. After fixation, embryos were stored in 1× PBS containing 0.02% sodium azide at 4°C until use.

For cryosectioning, embryos were washed twice with 1× PBS, then incubated in 15% (w/v) sucrose in 1× PBS at 4°C overnight or until fully submerged. This was followed by transfer to 30% (w/v) sucrose in 1× PBS at 4°C, again until tissues were fully infiltrated. Embryos were equilibrated in a 1:1 mixture of 30% sucrose and Tissue-Tek optimal cutting temperature (OCT) compound (VWR, 25608-930) for 2 hours at 4°C, then transferred into 100% OCT at 4°C for an additional 2 hours. Tissues were embedded in OCT using liquid nitrogen vapor and stored at –80°C. Cryosections were cut at a thickness of 12 µm using a Leica cryostat and mounted onto Superfrost Plus charged slides (VWR, 48311-703).

### Genotyping

Genomic DNA was isolated from limb bud tissue using the Extracta DNA Prep for PCR kit (Quantabio, 95091-025), following the manufacturer’s recommended protocol. To identify Elp1 conditional knockout (CKO) and control alleles, PCR was performed using the following primer pair: forward 5’-GCACCTTCACTCCTCAGCAT-3’ and reverse 5’-AGTAGGGCCAGGAGAGAACC-3’. Detection of the Wnt1-Cre transgene was carried out using primers 5’-GCCAATCTATCTGTGACGGC-3’ (forward) and 5’-CCTCTATCGAACAAGCATGCG-3’ (reverse). PCR reactions were assembled using DreamTaq Green PCR Master Mix (Thermo Fisher Scientific, K1082), prepared according to the manufacturer’s instructions.

### Protein Extraction, Digestion, and Mass Spectrometry

Tissue samples were lysed by heating in trifluoroacetic acid (TFA) at 73°C for approximately 5 minutes. Proteins were then neutralized by a 10-fold excess volume of 2 M Tris buffer containing 5 mM tris(2-carboxyethyl)phosphine (TCEP) and 10 mM chloroacetamide (CAA) to simultaneously reduce and alkylate cysteine residues. The samples were incubated at 90°C for 1 hour to ensure complete denaturation and reaction. Lysates were then diluted 6-fold with HPLC grade water containing 0.5 µg of sequencing-grade trypsin and incubated overnight at 37°C for proteolytic digestion.

Following digestion, peptides were acidified to a final concentration of 2% TFA and desalted using Evotips (Evosep) according to the manufacturer’s protocol. Peptides were separated using an Evosep One liquid chromatography system (EV-1000) employing an 88-minute gradient. Solvent A consisted of 0.1% formic acid in water, and Solvent B was 0.1% formic acid in acetonitrile. For each liquid chromatography mass spectrometry run, 20% of the peptide digest was used.

Mass spectrometry analysis was performed on a Thermo Scientific Q-Exactive HF-X Orbitrap operated in data-independent acquisition (DIA) mode. Full MS scans were acquired at a resolution of 120,000 with an AGC target of 3e6, a maximum injection time of 60 ms, and a scan range of 350–1650 m/z. DIA MS/MS spectra were collected at a resolution of 30,000 using an AGC target of 3e6, a fixed first mass of 200.0 m/z, and a stepped normalized collision energy (NCE) of 22.5, 25.0, and 27.5. DIA windows were set to 34.0 m/z.

Raw mass spectrometry data were processed in Spectronaut (Biognosys) using the directDIA workflow, which utilizes an in silico–predicted spectral library. Peptide and protein identification was performed against the UniProt mouse reference proteome (including isoforms), with the addition of a list of common laboratory contaminants. The search was set to trypsin ‘specific’ allowing up to 2 missed cleavages. Carbamidomethylation on cysteine was set as a fixed modification, while oxidation (M) and acetylation (Protein N-term) were set as variable modifications. Peptide and protein-level identifications were filtered to maintain a false discovery rate (FDR) below 1% and only proteins with two distinct peptides were evaluated further. The three most intense fragment ions of up to 3 different peptides per protein group were summed. The protein intensities were subjected to log2 transformation and subsequently normalized using variance stabilizing normalization. Imputation was performed using random draws from a Gaussian distribution centered around a minimal value, which in this case was the first percentile. Single-hit proteins were excluded from the analysis. T-tests were performed using a moderated t-test, after which multiple hypothesis testing (Benjamini & Hochberg, 1995) was applied to generate adjusted p-values. The data was filtered using a an adjusted p-value cut-off of <0.05.

### Immunohistochemistry

A hydrophobic barrier was drawn around each tissue section using an ImmEdge Pen (Vector Labs, H-4000) to contain reagents. Sections were rehydrated in 1× PBS for 5 minutes, followed by permeabilization in 0.5% Triton X-100 in 1× PBS for 10 minutes at room temperature. Blocking was carried out in PBS-Tx (1× PBS with 0.1% Triton X-100) containing 5% (w/v) bovine serum albumin (BSA, Fisher, BP9703100) for approximately 1 hour at room temperature. Sections were then rinsed once with PBS-Tx.

Primary antibodies, including goat anti-Gfra3 (R&D Systems, AF2645), rabbit anti-galanin (Abcam, ab32503), and rabbit anti-Chd1 (Proteintech, 20576-1-AP) were diluted in PBS-Tx with 1% (w/v) BSA and applied to the sections overnight at 4°C in a humidified chamber. After incubation, sections were washed four times with PBS-Tx for 5 minutes each at room temperature to remove unbound primary antibodies. Secondary antibodies, including donkey anti-goat 488 (Thermo, A-32814) and goat anti-rabbit 488 (Thermo, A-11008), were also diluted in PBS-Tx with 1% BSA, and applied for 1 hour at room temperature in a humidified chamber. Sections underwent three additional 5-minute rinses in PBS-Tx, followed by two final washes in 1× PBS for 5 minutes each.

Coverslips were mounted using DAPI Fluoromount-G Mounting Medium (Southern Biotech, 0100-20) and left to cure in the dark at room temperature overnight prior to imaging. Fluorescent signal was captured using a Zeiss LSM 800 confocal microscope with 10× and 20× air objectives. Imaging parameters, including laser power, gain, offset, digital zoom, and pinhole size (set to 1 airy unit), were held constant across Control and Elp1 CKO samples for comparative imaging. Image files (.CZI format) were processed using Zeiss Zen software (Blue edition 2.0), with histogram adjustments applied equally across experimental groups. Fluorescent signal was quantified using FIJI (Schindelin et al., 2012) measure application and statistical comparisons were performed using Prism (Graphpad).

---

**Supplemental Table 1.** Excel file listing identified proteins, log2-transformed fold changes, and p-values from proteomics experiment comparing E13.5 control and Elp1 CKO trigeminal ganglia.

